# Habitat transformation reshapes diversity and community structure of amphibians and reptiles in the Eastern Andes

**DOI:** 10.1101/2025.01.10.632384

**Authors:** Nelson Falcón-Espitia, Juan Camilo Ríos-Orjuela, Sebastian Perez-Rojas, Dennys Plazas-Cardona, Alejandra Arias-Escobar

**Affiliations:** Laboratorio de Biología Evolutiva de Vertebrados, Departamento de Ciencias Biológicas, Universidad de los Andes, 111711, Bogotá, Colombia; Grupo de Morfología y Ecología Evolutiva, Instituto de Ciencias Naturales, Universidad Nacional de Colombia, Sede Bogotá, 111321, Colombia; Independent researcher; SELVA: Investigación para la Conservación en el Neotrópico, 111311, Colombia

**Keywords:** Agroecosystems, anthropic disturbance, biodiversity conservation, habitat use, herpetofauna inventory

## Abstract

Land-use change is a major driver of biodiversity loss in tropical montane ecosystems, yet its effects on herpetofaunal community structure remain insufficiently understood at local scales. We evaluated patterns of diversity and community structure of amphibians and reptiles across a gradient of habitat transformation in southwestern Cundinamarca, Colombia. Field surveys conducted between 2015 and 2022 using visual encounter surveys recorded 57 species and 608 individuals. Forest habitats supported the highest species richness and diversity, whereas coffee crops and open areas exhibited reduced richness and simplified assemblages. Rank–abundance patterns revealed greater dominance in open habitats and higher evenness in forests, consistent with environmental filtering processes. Amphibians were more strongly associated with forest environments, whereas reptiles showed broader habitat tolerance and higher representation in disturbed habitats. In addition, we compiled a comprehensive regional inventory based on field data, biological collections, and biodiversity databases, documenting 86 species of herpetofauna. Overall, our results show patterns consistent with environmental filtering, where habitat transformation is associated with increased dominance of generalist species and simplified community structure. These findings highlight the importance of conserving forest remnants and maintaining heterogeneous landscapes to support herpetofaunal diversity in the Colombian Andes.

## Introduction

In Colombia, Andean ecosystems have undergone extensive transformation, particularly in Andean and sub-Andean forests, which concentrate the highest levels of productive activity in the country and, consequently, strong anthropogenic pressures mediated by ranching, agriculture, infrastructure development and mining (Correa-Ayram et al. 2020, Rubiano et al. 2026). The Andean region harbors approximately 53% of the country’s total herpetofauna (Romero et al. 2009), including a high number of endemic species (Bernal and Lynch 2008, Armesto and Señaris 2017). At a broader scale, the tropical Andes are recognized as a global biodiversity hotspot (Myers et al. 2000, Vasconcelos et al. 2019), characterized by exceptional levels of endemism and a high concentration of threatened species (Tobar-Suárez et al. 2022, IUCN 2024, Vásquez-Restrepo and García-Cobos 2026). Additionally, a substantial proportion of these species occurs outside protected areas, increasing their vulnerability to ongoing land-use change, particularly in human-dominated landscapes (Nori et al. 2015, Bax and Francesconi 2019, Vásquez-Restrepo and García-Cobos 2026).

One of the main drivers shaping amphibian and reptile assemblages in tropical landscapes is land-use transformation associated with agriculture, livestock, and urban expansion (Ghosh and Basu 2020, Iglesias-Carrasco et al. 2023). These processes lead to habitat loss, degradation, and fragmentation, resulting in heterogeneous landscapes composed of remnant forest patches embedded within human-modified matrices (Thompson et al. 2015, Cervantes-López and Morante-Filho 2024). Such transformations affect not only species richness, but also the composition and internal structure of ecological assemblages, altering patterns of dominance, evenness, and species interactions (Ernst et al. 2006, Gardner et al. 2007). As a result, disturbed habitats are often characterized by simplified assemblages dominated by a reduced number of species. However, beyond changes in species richness, how habitat transformation reshapes the internal structure of assemblages, including patterns of dominance and evenness, remains less clearly understood.

Differences in species responses to habitat transformation are strongly mediated by physiological and ecological traits. Amphibians, due to their permeable skin, reproductive dependence on moisture, and narrow thermal tolerances, are generally more sensitive to environmental changes (Wanger et al. 2010, Palacios et al. 2013, Méndez-Narváez 2014, Sankararaman and Miller 2024). In contrast, reptiles tend to exhibit broader tolerance to thermal and hydric variation, allowing them to persist or even increase in abundance in disturbed environments (Thompson et al. 2015, Cordier et al. 2021, Veloza and Urbina-Cardona 2025). These contrasting responses are consistent with environmental filtering processes, providing a mechanistic framework to understand how habitat transformation restructures ecological assemblages through shifts in dominance and species composition (Brüning et al. 2018, Zabala-Forero and Urbina-Cardona 2021, López-Bedoya et al. 2022). Understanding these patterns is particularly relevant in tropical montane systems, where environmental gradients and land-use change interact to shape biodiversity.

In heterogeneous landscapes, agroecosystems may play a significant role in conserving biodiversity. Some agricultural systems, such as shaded coffee crops, have been proposed as biodiversity-friendly land uses capable of maintaining a subset of native species while supporting economic production (Perfecto and Vandermeer 2008, Céspedes and Bayly 2019, González et al. 2020, Ríos-Orjuela et al. 2024). These systems can function as intermediate habitats or dispersal matrices between forest remnants. However, despite increasing evidence of the effects of land-use change, there is still limited understanding of how multiple dimensions of diversity and community structure respond simultaneously across heterogeneous landscapes at local scales, particularly in regions with complex mosaics of land use (Brüning et al. 2018, Agudelo-Hz et al. 2019, Roach et al. 2020, Zabala-Forero and Urbina-Cardona 2021).

Here, we evaluated how habitat transformation shapes diversity and community structure of amphibians and reptiles across a heterogeneous landscape in the southwestern Cundinamarca region. Specifically, we asked whether changes in habitat structure are associated with shifts in species richness, dominance patterns, and community composition across forest, agricultural, and open habitats. Based on environmental filtering theory, we expected that: (1) species richness would decline from forest to more transformed habitats; (2) community structure would shift towards increased dominance and reduced evenness in disturbed habitats; and (3) responses would differ between taxonomic groups, with amphibians more strongly associated with forest environments and reptiles showing broader tolerance to disturbance. These predictions reflect the expectation that habitat transformation selectively favors generalist species while restricting taxa with more specialized ecological requirements.

Rather than directly testing causal mechanisms, our aim is to provide a standardized, coverage-based characterization of diversity and community structure across habitats, generating a robust baseline for understanding how habitat transformation reshapes assemblages. In addition to these analyses, we compiled historical and current occurrence records to generate a comprehensive species inventory for the region, providing a broader biogeographic context for interpreting observed patterns of diversity and community structure.

## Methods

### Study area

We conducted fieldwork in three municipalities of the southwestern Cundinamarca region (Tibacuy, Nilo, and Viotá), located in the Eastern Cordillera of the Colombian Andes, between 645 and 1840 m elevation (Fig. 1). The study area comprises heterogeneous landscapes dominated by secondary Andean forest and agroecosystems within a mosaic of forest remnants, shaded coffee crops and open areas used for livestock farming (Peña-Torres 2016).

**Figure 1.**
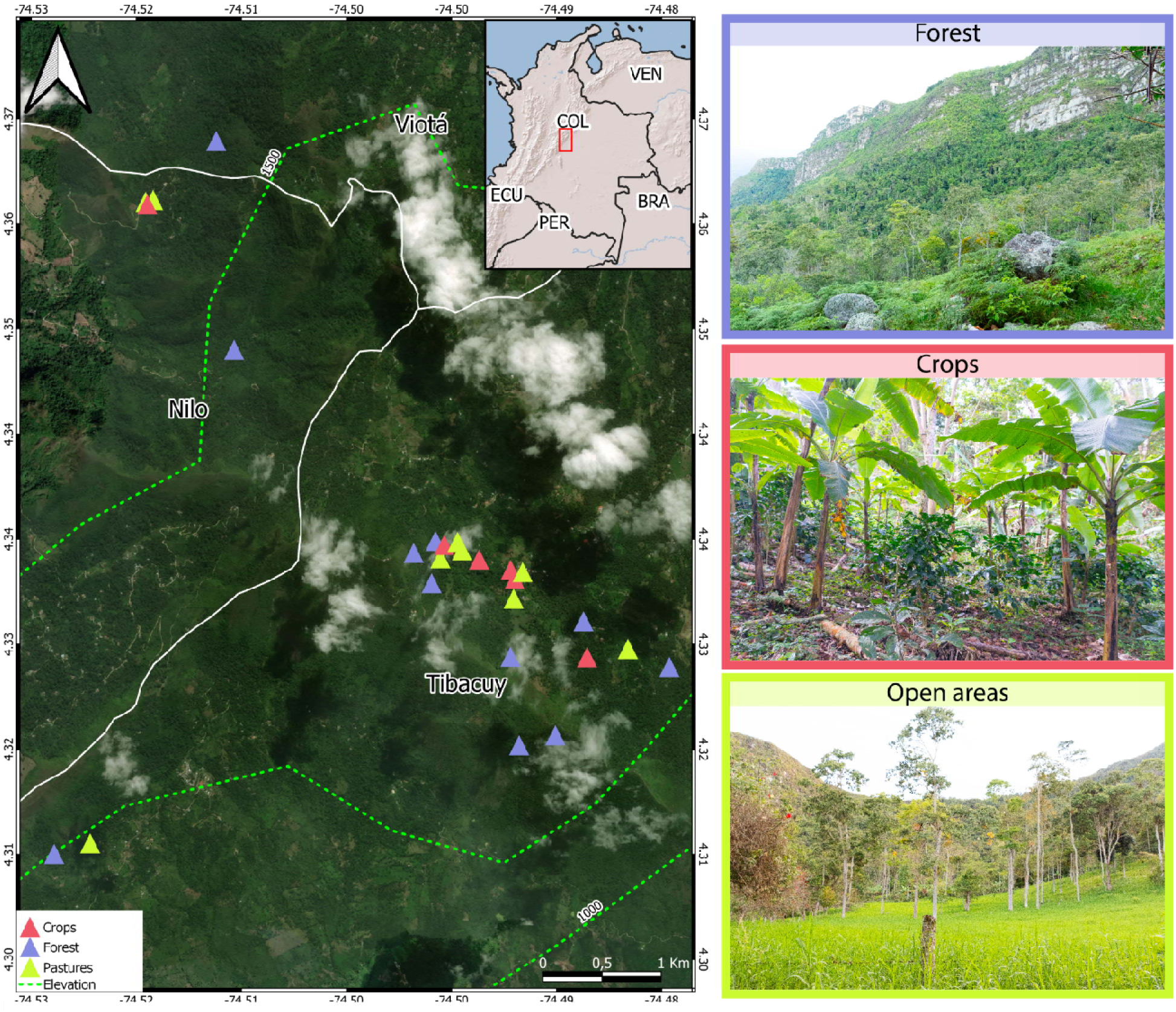
Study area and sampling design across a heterogeneous landscape in the southwestern Cundinamarca region in the Eastern Cordillera. Herpetofaunal sampling was conducted across a heterogeneous landscape dominated by forest, crops, and open areas along an elevational gradient, providing the spatial framework for comparing community structure among habitats. The map shows sampling localities in the municipalities of Tibacuy, Nilo, and Viotá (Cundinamarca, Colombia), with points colored by habitat type: forest (blue), crops (red), and open areas (green). Elevational contours (dashed lines) indicate the altitudinal gradient across the study area (645–1840 m). The inset locates the study region within Colombia. Representative photographs illustrate the three habitat types surveyed: forest (top), shaded coffee crops (middle), and open areas (bottom).

According to Holdridge’s life zone classification (1967) the region includes low-montane rainforest and low-montane dry forest zones characterized by predominantly humid mountain climate in combination with local dry enclaves, typical of the middle lands of the Magdalena basin. Sampling was conducted across three habitat types: secondary forest (11 localities), with scattered trees over five meters high, where advanced plant regeneration processes are evident; shaded coffee crops (6 localities) and open areas (9 localities) used for permanent livestock. Geographic coordinates, elevation, sampling period, and number of transects per locality are provided in Supplementary Material 1.

### Field sampling and sampling effort

Herpetofauna surveys were conducted in two periods: between 2015 and 2019 and between June 2021 and July 2022, covering the rainy and dry seasons in the region. During the sampling phase, we employed time-constrained Visual Encounter Surveys (VES), following standard protocols for amphibian and reptile inventories (Villarreal et al. 2006). Sampling consisted of active visual and auditory surveys within representative microhabitats including vegetation, leaf litter, streams, and ponds, up to a maximum height of 5 m above ground (Cortés et al. 2008). Auditory detections were estimated by a single observer to ensure consistency and represent approximate counts of calling individuals.

Each sampling unit consisted of a four-hour search conducted by five observers. Surveys were performed during both diurnal (8:00-12:00) and nocturnal (18:00–22:00 h) activity periods. In total, 53 sampling events were conducted across the study area between 2015 and 2022, corresponding to a total sampling effort of 1060 person-hours. Sampling effort was broadly comparable across habitats, although it varied slightly among them, with 500 person-hours in forest habitats, 340 person-hours in crops, and 220 person-hours in open areas. Survey sites were selected within representative areas of each habitat type. Not all localities were visited in every sampling period; however, each sampling event was treated as a sampling unit within habitat categories, as analyses were designed to compare community patterns among habitats rather than to infer independent population-level processes. The distribution of sampling localities, periods of visitation, and number of transects per site are detailed in Supplementary Material 1.

### Community analyses

Habitat types were defined based on vegetation structure and land use following a simplified classification scheme. Forest sites correspond to areas with continuous or semi-continuous tree cover and advanced vegetation structure. Coffee crops include shaded or semi-shaded agricultural systems with varying management intensity. Open areas comprise pasture-dominated landscapes, including grasslands with sparse or no tree cover. These categories represent a gradient of habitat transformation from less disturbed (forest) to highly modified environments (open areas). A detailed description of vegetation cover units is provided in Ríos-Orjuela et al. (2024).

To evaluate species richness at standardized sampling efforts and predict species richness beyond our observed sample size, we assessed sampling completeness using coverage curves for the entire community and by habitats, using rarefaction and extrapolation analyses based on Hill numbers with 1000 bootstrap resampling to estimate 95% confidence intervals (CI). We quantified diversity as Hill numbers representing species richness (q0), Shannon diversity (q1), and Simpson diversity (q2) using the *iNEXT* package (Chao et al. 2014, Hsieh et al. 2016).

To evaluate patterns of species dominance and evenness among habitats, we calculated rank–abundance curves using abundance data grouped by habitat type. Amphibians and reptiles were analyzed separately to better illustrate differences in community structure between taxonomic groups. We log-transformed abundance values to improve visualization of rank distributions. In addition, rank-abundance plots were generated using *ggplot2* (Wickham 2009), and annotated with *ggrepel* to identify dominant species within each habitat.

Finally, to assess potential spatial autocorrelation in species richness among sampling localities, Moran’s I statistic was calculated using species richness per locality and corresponding geographic coordinates, defining spatial weights and using a k-nearest neighbor approach with the *spdep* package (Bivand et al. 2026).

Given the descriptive nature of this study, comparisons among habitats are based on coverage-standardized diversity estimates and their associated confidence intervals, rather than formal hypothesis-testing approaches. All ecological analyses were conducted exclusively using field survey data collected between 2015–2019 and 2021–2022. All analyses were performed in R version 4.1.0. (R Core Team 2023).

### Regional diversity compilation

To compile a comprehensive species inventory for the study region, field records were complemented with additional occurrence data obtained from the Biodiversity Information System of Colombia (SiB Colombia) and biological collections. Records were filtered to include preserved specimens and verified institutional observations to minimize potential identification errors. We also reviewed specimen records from the Instituto de Ciencias Naturales (ICN) at Universidad Nacional de Colombia and the amphibian and reptile collections of the Instituto de Investigación de Recursos Biológicos Alexander von Humboldt (IAvH) for the municipalities of Tibacuy, Nilo, and Viotá (Cundinamarca). These supplementary records were used exclusively to compile the regional species list and were not included in ecological analyses. Finally, we followed Frost (2024) and Uetz et al. (2024) as taxonomy classification systems for amphibians and reptiles, respectively.

## Results

### Field surveys

Field surveys conducted between 2015-2019 and 2021-2022 recorded a total of 57 species, representing a substantial component of the known regional diversity (Supplementary Material 2). Across all sampling events, a total of 608 individuals were recorded, including 391 amphibians and 217 reptiles. The most abundant amphibian species were *Boana platanera* (62 individuals), *Engystomops pustulosus* (60), and *Pristimantis taeniatus* (55). Among reptiles, the most abundant species were *Gonatodes albogularis* (53 individuals), *Cnemidophorus lemniscatus* (33), and *Hemidactylus frenatus* (17).

Species richness varied among habitat types, with forest supporting the highest richness (45 species), followed by open areas (34) and coffee crops (14). In terms of abundance, forests also contained the greatest number of individuals (225 records), followed by open areas (207), while coffee crops supported considerably fewer individuals (176).

### Sampling completeness

Rarefaction and extrapolation analyses showed that sampling covered a large proportion of the expected diversity of both amphibians and reptiles (Fig. 2a). Sample coverage exceeded 95% for both taxonomic groups (Fig. 2b). Coverage-based rarefaction curves approached asymptotic levels, with amphibians reaching asymptotes at lower sample sizes than reptiles.

**Figure 2.**
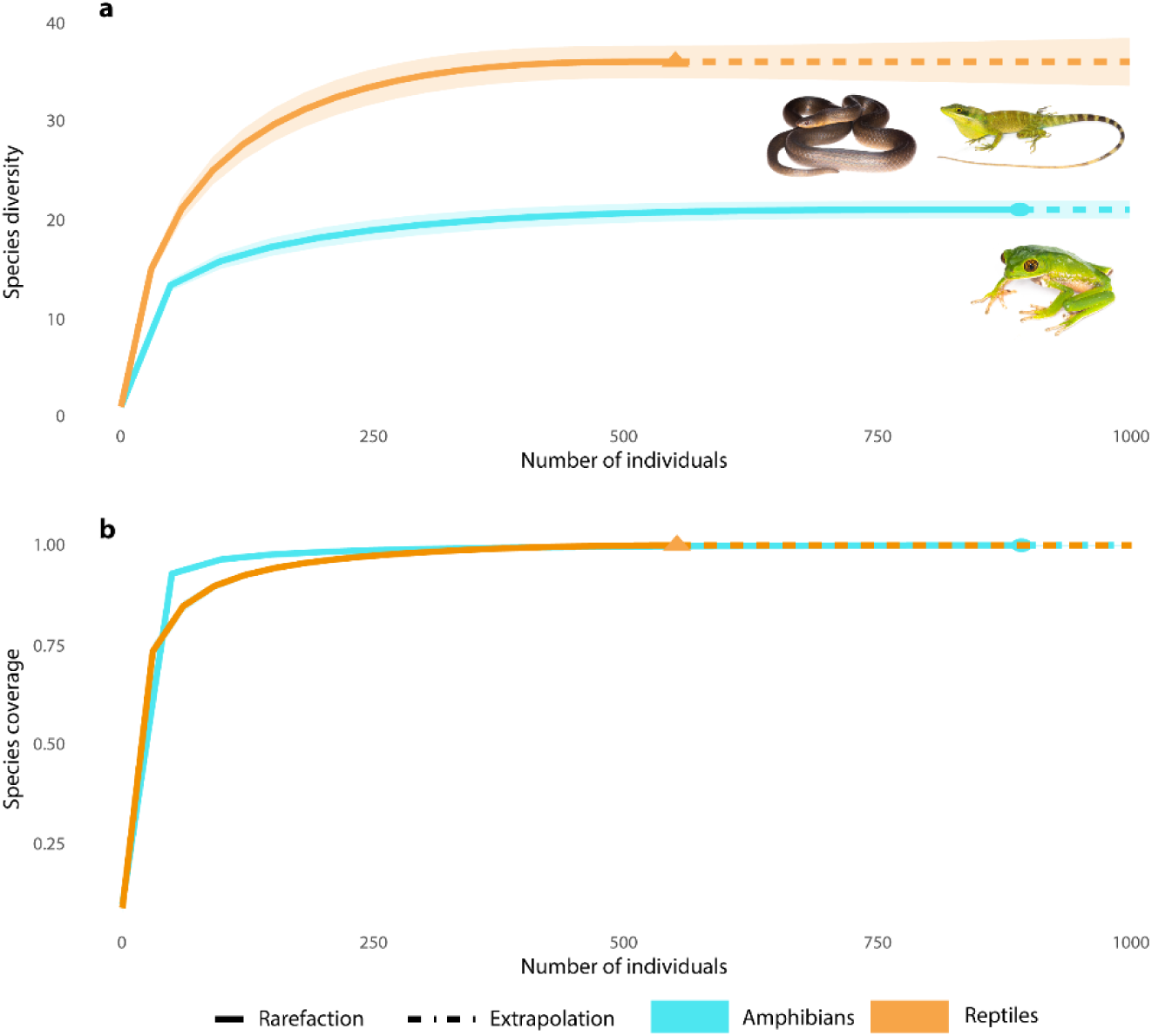
Sampling captured most of the expected diversity, with higher species richness in reptiles than amphibians. Sampling achieved high completeness for both taxonomic groups, and reptiles consistently exhibited higher species richness than amphibians across the full range of observed and extrapolated sample sizes. (a) Rarefaction (solid lines) and extrapolation (dashed lines) curves show the relationship between species diversity (Hill number q0) and the number of individuals sampled for amphibians (blue) and reptiles (orange). Symbols indicate the observed sample size, and shaded areas represent 95% confidence intervals based on 1,000 bootstrap replicates. (b) Sample coverage curves show that sampling completeness rapidly approached asymptotic values (>95%) for both groups.

Spatial autocorrelation analysis using Moran’s I revealed no significant spatial structure in species richness across sampling localities (Moran’s I = −0.0147, expected I = −0.040, p = 0.385), indicating that observed richness patterns of the herpetofauna community were not spatially structured.

### Diversity across habitats

Coverage-based rarefaction and extrapolation analyses showed differences in diversity among habitats (Fig. 3). Forest exhibited the highest estimated diversity across all Hill numbers with estimated species richness (q0) in forests reaching 57.6 species (95% CI: 33.9–81.3), compared to 35.5 species in crops (95% CI: 27.3–43.6) and 23.3 species in open areas (95% CI: 14.4–32.2). Shannon diversity (q1) showed a similar pattern, with forests exhibiting the highest effective number of species (21.7; 95% CI: 17.4–26.1), followed by crops (16.6; 95% CI: 14.0–19.2) and open areas (15.2; 95% CI: 12.6–17.8). Simpson diversity (q2) was similar between forests (11.4; 95% CI: 8.8–13.9) and open areas (12.2; 95% CI: 10.2–14.2), whereas crops exhibited lower dominance-adjusted diversity (8.8; 95% CI: 6.9–10.6). Confidence intervals for species richness (q0) did not overlap between forest and open areas, whereas estimates for other diversity metrics showed partial overlap among habitats.

**Figure 3.**
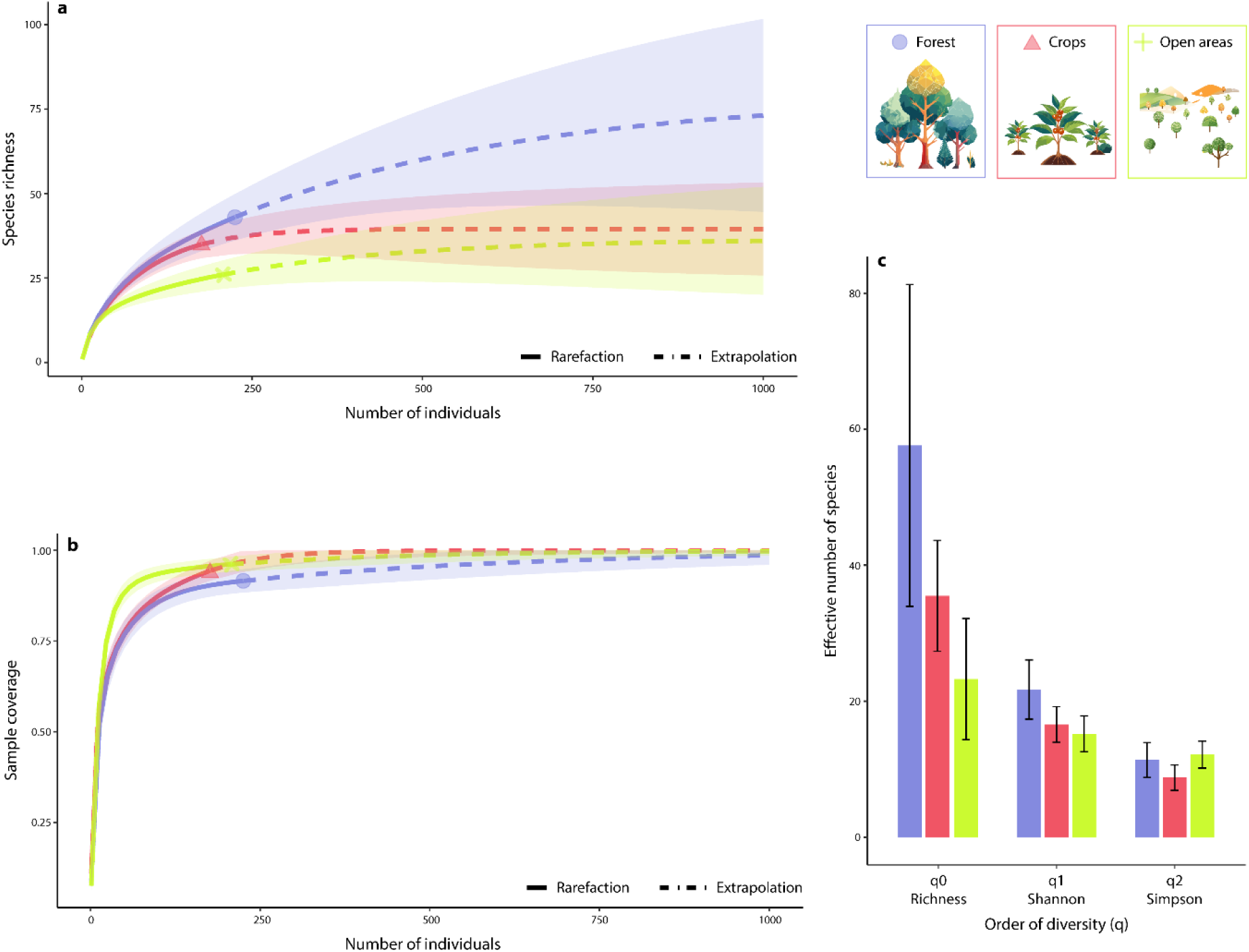
Forest habitats support higher diversity, whereas transformed habitats show reduced richness and simplified assemblages. Species diversity declined from forest to crops and open areas, suggesting that habitat transformation reduces richness and simplifies community structure across the landscape. (a) Coverage-based rarefaction (solid lines) and extrapolation (dashed lines) curves of species richness (Hill number q0) as a function of the number of individuals sampled for forest (blue), crops (red), and open areas (green). Symbols indicate observed sample sizes, and shaded regions represent 95% confidence intervals based on 1,000 bootstrap replicates. (b) Sample coverage curves show that sampling completeness was similarly high across habitats, allowing robust comparisons of diversity patterns. (c) Effective diversity (Hill numbers q0, q1, q2) by habitat type. Forest exhibits the highest richness and Shannon diversity, whereas crops and open areas show reduced diversity. Differences in Simpson diversity (q2) suggest comparable dominance structure between forest and open areas, but reduced diversity in crops. Error bars represent 95% confidence intervals.

When analyzed separately, amphibians and reptiles also showed distinct diversity patterns across habitats. Amphibian richness was highest in forests and declined in crops and open areas. Assemblages in open areas showed relatively higher evenness compared to other habitats. Reptiles followed a similar gradient in species richness, with forests supporting the highest diversity, while open areas showed lower diversity and stronger dominance patterns. In crops, reptile assemblages exhibited relatively even distributions of abundances. Detailed diversity estimates for amphibians and reptiles across habitats are provided in Supplementary Material 3.

### Community structure across habitats

Rank–abundance distributions revealed differences in species dominance and community structure among habitats (Fig. 4). Forest assemblages showed longer tails in rank distributions, with a greater number of low-abundance species.

**Figure 4.**
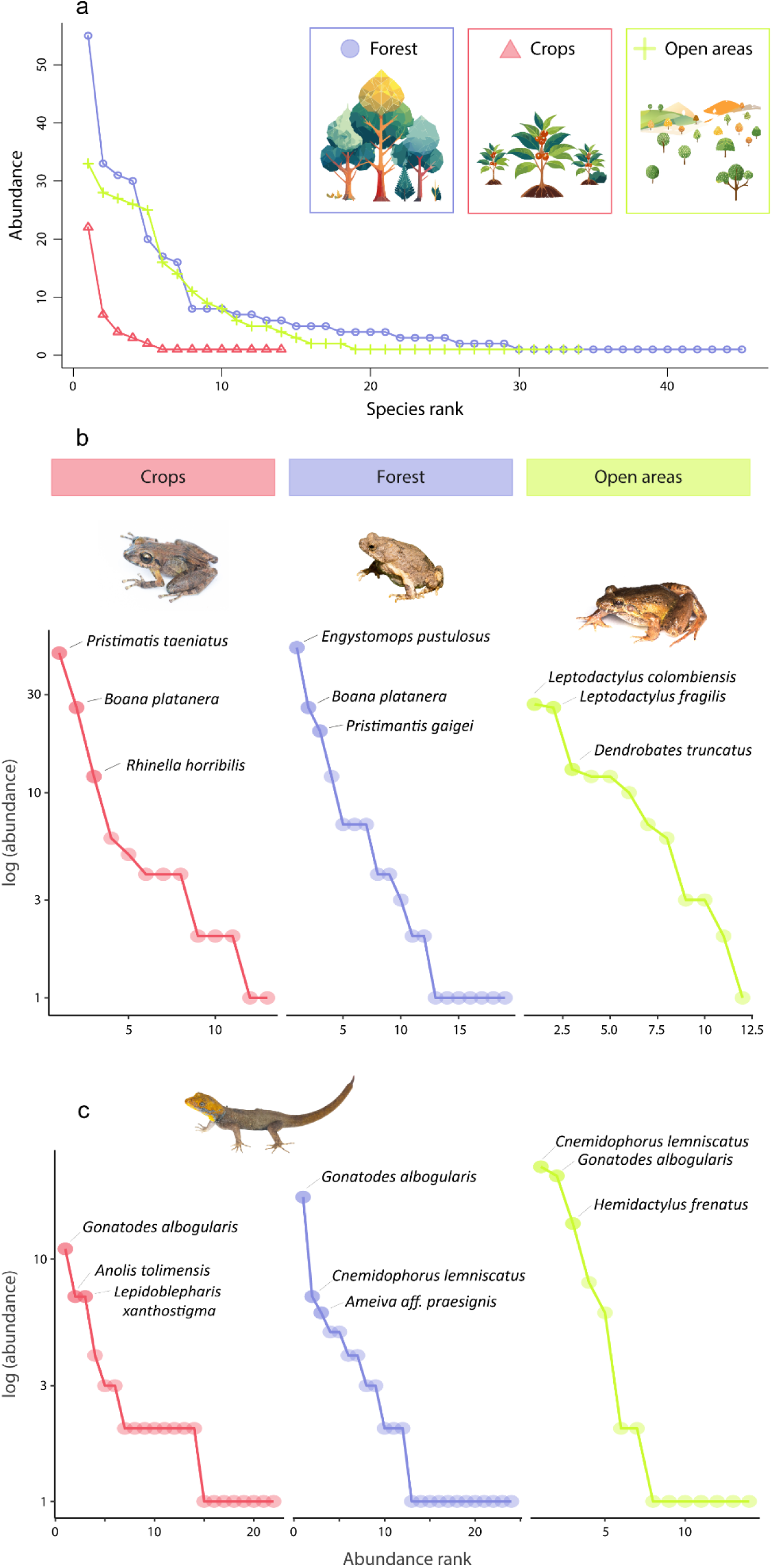
Habitat transformation increases dominance and simplifies community structure, favoring a few abundant species in disturbed habitats. Community structure shifts from more even assemblages in forests to dominance by a few species in transformed habitats, suggesting that habitat modification favors generalist species and reduces community complexity. (a) Rank–abundance curves for all species pooled by habitat type. Forest (blue) shows a longer tail and more gradual decline in abundance, showing higher evenness and greater representation of rare species. In contrast, crops (red) and open areas (green) exhibit steeper declines, reflecting stronger dominance by a few species. (b) Rank–abundance curves for amphibians by habitat type (log-transformed abundance). Forest assemblages display greater richness and evenness, whereas crops and open areas are dominated by a reduced number of species. Representative dominant species are labeled for each habitat. (c) Rank–abundance curves for reptiles by habitat type (log-transformed abundance). Open areas show strong dominance by a few widespread species (e.g., *Cnemidophorus lemniscatus, Gonatodes albogularis,* and *Hemidactylus frenatus*), whereas forest assemblages exhibit a more even distribution of abundances across species. Species labels highlight the most abundant taxa in each habitat.

Forest showed higher richness and a more gradual decline in amphibian species abundance, whereas coffee crops and open areas were characterized by fewer species and stronger dominance by a small subset of taxa (Fig. 4b). Dominant amphibian species varied among habitats, including *Engystomops pustulosus*, *Boana platanera*, and *Pristimantis* species. Open areas were dominated by a few highly abundant species such as *Cnemidophorus lemniscatus, Gonatodes albogularis*, and *Hemidactylus frenatus*, whereas forest sites contained a broader distribution of species with lower individual abundances (Fig. 4c).

Overall, these results describe clear differences in community structure across habitat types, with forests showing higher species richness and more even assemblages, and open areas characterized by greater dominance of fewer species.

### Regional herpetofauna checklist

A total of 86 species of herpetofauna have been recorded in the study region based on our field surveys, biological collections, and biodiversity records in databases (Supplementary Material 2). Among amphibians, the order Anura is the richest group with 32 of 35 species recorded. The most speciose families were Hylidae (8 species), followed by Strabomantidae and Dendrobatidae (5 species each). Caecilians (Gymnophiona) were represented by two species whereas there are reports of a single species of salamander (Caudata) in the area. Regarding reptiles, 51 species have been recorded primarily represented by the order Squamata (49); the most diverse families were Colubridae (23 species) and Anolidae (6 species). The orders Crocodylia and Testudines were represented by a single species each (*Caiman crocodilus* and *Kinosternon* sp.). Two invasive species were recorded in the region: the North American bullfrog *Aquarana catesbeiana* and the South Asian house gecko *Hemidactylus frenatus*.

## Discussion

### Drivers of community structure across habitats

Habitat transformation is a major driver of biodiversity patterns in tropical landscapes, affecting both species richness and the internal structure of ecological assemblages (López-Bedoya et al. 2022, Cervantes-López and Morante-Filho 2024). In this study, herpetofaunal assemblages were strongly structured by habitat type, with secondary forests supporting the highest diversity, while agricultural and open areas hosted distinct but simplified assemblages. Despite reduced diversity in modified habitats, a substantial portion of the regional fauna persisted across habitat types, indicating that different components of the landscape contribute unevenly but complementarily to overall diversity.

Although sampling units were spatially connected, the absence of spatial autocorrelation and the consistent differentiation among habitats suggest that environmental filtering, rather than spatial dependence, plays a central role in structuring community patterns. Under this framework, habitat transformation is associated with assemblages dominated by generalist species and reduced representation of taxa with more specialized ecological requirements.

At a regional scale, the southwestern Cundinamarca region harbors a relatively diverse herpetofauna, based on field surveys and compiled records (Lynch 1982, Ramírez 2017, Acosta-Galvis et al. 2020, López-Perilla et al. 2023). Amphibians and reptiles were represented by several families with multiple species, including Colubridae, Anolidae, Hylidae and Strabomantidae, and sampling coverage indicates that the assemblages documented here provide a representative characterization of the local herpetofauna. This diversity reflects the convergence of forest remnants and human-modified habitats across the transition between lowland tropical forests and montane environments, which contributes to shaping regional biodiversity patterns (Brüning et al. 2018, Zabala-Forero and Urbina-Cardona 2021, López-Bedoya et al. 2022).

### Diversity patterns across habitat transformation

Clear differences in diversity among habitats reflect variation in structural complexity and microclimatic stability (Cortés-Gómez et al. 2013, Roach et al. 2020, Cervantes-López and Morante-Filho 2024), which promote the persistence of species with specialized ecological requirements and complex life histories (Veloza and Urbina-Cardona 2025). Forest habitats supported the highest diversity across all metrics, whereas crops and open areas exhibited reduced diversity and altered community structure.

These patterns are consistent with previous studies showing that habitat modification reduces species richness and modifies community structure in tropical systems (Roach et al. 2020, Galindo-Uribe et al. 2022, Veloza and Urbina-Cardona 2025). In particular, the non-overlapping confidence intervals observed between forest and open areas for species richness highlight a marked difference in diversity between less disturbed and highly transformed habitats, while partial overlap in other diversity metrics suggests more subtle changes in dominance and evenness.

### Contrasting responses of amphibians and reptiles

Differences between amphibians and reptiles may reflect contrasting physiological constraints and ecological strategies. Amphibians were more strongly associated with forest habitats, likely due to their dependence on humid and thermally stable microclimates, which increase the risk of desiccation and reduce reproductive success in disturbed environments (Wanger et al. 2010, Palacios et al. 2013, Méndez-Narváez 2014, Cervantes-López and Morante-Filho 2024).

In contrast, reptiles exhibited broader habitat occupancy, consistent with their higher tolerance to thermal and hydric variation (Méndez-Narváez 2014, Thompson et al. 2015, Cordier et al. 2021, Cervantes-López and Morante-Filho 2024). These differences support the expectation that taxonomic groups respond differently to habitat transformation depending on their physiological and ecological traits.

### Community dominance and structural simplification

Patterns of dominance varied across habitats, with open areas characterized by assemblages dominated by a small number of widespread species (Fig. 4), a pattern frequently associated with ecological disturbance and habitat simplification (Flynn et al. 2009, Herrera-Montes and Brokaw 2010) and previously reported in the region (Veloza and Urbina-Cardona 2025). In contrast, forest assemblages exhibited a more even distribution of abundances and a greater representation of low-frequency species.

The distribution of species across habitats reflects differences in ecological strategies. Generalist species occurred consistently across all habitats (Supplementary Material 2) and tend to dominate local assemblages, indicating high tolerance to environmental variation (Brüning et al. 2018, Zabala-Forero and Urbina-Cardona 2021, Ríos-Orjuela et al. 2024). In contrast, forest habitats harbored species with more restricted ecological requirements, particularly amphibians dependent on stable microclimatic conditions (Galindo-Uribe et al. 2022). These patterns are consistent with a shift toward generalist-dominated assemblages in transformed habitats.

### Role of agroecosystems in heterogeneous landscapes

Agricultural systems may play an important role within heterogeneous landscapes (Galindo-Uribe et al. 2022, Cervantes-López et al. 2025). In the study area, coffee crops hosted several species also present in forest habitats, indicating that these environments can function as secondary habitats or dispersal matrices (Ríos-Orjuela et al. 2024). Agroecosystems therefore may contribute to biodiversity conservation when embedded within heterogeneous landscapes (Roach et al. 2020, Zabala-Forero and Urbina-Cardona 2021, Ríos-Orjuela et al. 2024, Cervantes-López et al. 2025, Pinzón et al. 2025). However, these habitats do not replace forest ecosystems, as they primarily support disturbance-tolerant species and retain only a subset of the regional fauna. Additional dimensions of diversity, such as functional and phylogenetic diversity should be taken into account to better understand the herpetofaunal assemble dynamics and the role of agroecosystems in conserving community structure.

### Broader implications: land-use change and biotic homogenization

The observed patterns should be interpreted within the broader context of land-use change in the Colombian Andes, where deforestation and agricultural expansion have driven habitat fragmentation (Armenteras et al. 2011, Rodríguez Eraso et al. 2013, Vanegas-Cubillos et al. 2022). These processes tend to favor ecological generalists while reducing the persistence of species with specialized requirements, potentially leading to biotic homogenization and changes in ecosystem functioning (Valencia-Aguilar et al. 2013, Thompson et al. 2015).

In this context, the presence of invasive species such as *Aquarana catesbeiana* and *Hemidactylus frenatus*, which have been in the region for decades (Rueda-Almonacid 1999, Caicedo-Portilla and Dulcey-Cala 2011, Pérez-Rojas et al. 2026), may further reinforce homogenization processes by favoring disturbance-tolerant assemblages (Rueda-Almonacid 1999, Corporación Autónoma Regional de Cundinamarca CAR 2018, Galindo-Uribe et al. 2022) in a region with one of the highest rates of endemism in amphibians and reptiles (Tobar-Suárez et al. 2022, Díaz-Ricaurte et al. 2025, Vásquez-Restrepo and García-Cobos 2026).

### Limitations and future directions

Our results should be interpreted considering some limitations. Although sampling coverage was high, detectability may vary among taxa and habitats, particularly between amphibians and reptiles. In addition, the elevational gradient encompassed in the study may partially confound habitat effects, as environmental conditions covary with both elevation and land use. Finally, given the descriptive nature of our analyses, the patterns documented here should be interpreted as indicative of underlying ecological processes rather than direct tests of causal mechanisms.

### Regional inventory and taxonomic considerations

In addition to providing ecological insights, this study contributes to improving the knowledge of the regional herpetofauna by presenting the first compiled species list for the southwestern Cundinamarca region in Colombia. Several recorded species represent expected occurrences based on their known distributions in nearby localities but had not previously been documented in detail for the study area. For instance, species such as *Leptodactylus fragilis*, *Rhinella humboldti*, *Cnemidophorus lemniscatus*, and *Bothrops asper* are widely distributed in the Magdalena River valley and surrounding regions, but their occurrence in the surveyed area contributes to refining the regional inventory. These records provide a more robust baseline for future ecological and conservation studies.

There are some records for both amphibians and reptiles worth discussing. Based on currently available morphological information for the species in the region, we were unable to determine one species of *Anolis*. Preliminary observations suggest affinity with *Anolis granuliceps*, although further integrative taxonomic analyses will be required to clarify its identity. The presence of potentially undescribed or poorly characterized taxa highlights the still incomplete knowledge of herpetofauna diversity in the Colombian Andes. Recent studies in the region have revealed several previously unrecognized lineages such as *Anolis tequendama* (Moreno-Arias et al. 2023), *Bolitoglossa muisca* (López-Perilla et al. 2023) and *Hyloxalus arliensis* (Acosta-Galvis et al. 2020). Likewise, recent systematic studies in *Scinax* suggest that populations currently assigned to this widespread species complexes may correspond to distinct evolutionary lineages (Guarnizo et al. 2015, Araujo-Vieira et al. 2020, 2023).

There are also several relevant historical records in the municipalities studied, especially toward higher altitudes. Species such as *Atelopus subornatus* and *Hyloxalus ruizi* were observed in the area surrounding El Alto de San Miguel by Lynch (1982). Both species are critically endangered (IUCN 2024), and besides the southern records of *A. subornatus* in Tolima (Enciso-Calle et al. 2017), there are no recent records in the area, and we did not register them in this study. Other species currently threatened in this area worth mentioning are *Hyloxalus vergeli, Pristimantis uisae* and *P. renjiforum* (Supplementary Material 2).

### Conservation implications

Our results underscore the importance of maintaining forest remnants and promoting landscape heterogeneity in the conservation of amphibians and reptiles in the Colombian Andes. Given that habitat transformation is associated with reduced representation of forest-dependent species, continued habitat loss is likely to disproportionately affect taxa with specialized ecological requirements.

Conservation strategies should prioritize the protection of remaining forest patches, the maintenance of ecological connectivity, and the implementation of land-use practices that promote structurally heterogeneous landscapes. Long-term monitoring programs will be essential to evaluate how ongoing environmental changes, including land-use transformation and climate change, affect the distribution and persistence of amphibian and reptile assemblages in Andean ecosystems (López-Bedoya et al. 2022, Cervantes-López and Morante-Filho 2024, Ríos-Orjuela et al. 2024).

## Acknowledgements

We thank the Universidad Nacional de Colombia for the permission to conduct this study (Resolución 255 de 2014 granted by ANLA). We thank the Asociación Colombiana de Herpetología for funding though the Botas al Campo grant (2019) and APRENAT for their logistical support. We also acknowledge the cooperation of various individuals and organizations, including the Castillo Urrego family, Bernal Méndez family, Ramos family, Maria Rosa Gaona de Chacón, Mayelly Liévano, Gabriel Garzón, Fray José María Sepúlveda, the Province of Nuestra Señora de la Gracia de Colombia and the herpetology group of Universidad Nacional de Colombia (Herpetos UN), for their hospitality, collaboration throughout the field phase and for their valuable contributions in the development of this project. To Nataly Casas, Carolina Martínez and Juan Diego Rodríguez for their help during the fieldwork phase and species identification. Also, we appreciate the ongoing efforts of different communities in Tibacuy, Nilo and Viotá municipalities in conserving their biocultural heritage. Finally, the authors are grateful to Dr. Liliana Saboyá and an anonymous reviewer for their suggestions, which improved the quality of this manuscript.

